# Protective Effects of Minocycline and Acetazolamide on Visual Function in Simulated Microgravity Rats

**DOI:** 10.1101/2025.04.06.647441

**Authors:** Liu Jinshuo, Yan Naiqin, Mao Yingyan, Xin Chen, Mou Dapeng, Wang Ningli, Zhu Siquan

## Abstract

This study established a tail-suspension rat model to investigate the effects of simulated microgravity on intraocular biomechanical homeostasis and visual function, while evaluating the protective roles of acetazolamide (AZE) and minocycline (MINO). Forty-eight rats were divided into ground control (CTRL, n=12) and simulated microgravity groups (n=36), with the latter further categorized by treatment (PBS, AZE, or MINO) and exposure duration (2 or 4 weeks; n=6/group). Assessments included intraocular pressure (IOP), intracranial pressure (ICP), translaminar cribrosa pressure difference (TLCPD), electroretinography (ERG), optomotor response (OMR), and optical coherence tomography (OCT), supplemented by immunofluorescence staining and Sholl analysis. Simulated microgravity significantly elevated IOP and ICP, altered TLCPD, and induced neuroimmune activation, leading to ERG and OMR impairments. AZE effectively reduced IOP and ICP, mitigating mechanical stress, whereas MINO suppressed microglial and astrocytic activation, attenuating retinal neurodegeneration. ERG and OMR demonstrated that MINO restored photopic negative response (PhNR) amplitudes near control levels by 4 weeks. OCT revealed that both AZE and MINO inhibited retinal thinning, particularly in the outer nuclear layer (ONL). Immunofluorescence confirmed MINO’s suppression of microglial activation and morphological changes. These findings suggest that AZE preserves visual function by modulating intraocular pressure dynamics, while MINO exerts neuroprotection by mitigating neuroinflammation, offering potential dual therapeutic strategies for spaceflight-associated visual impairment.

## Introduction

Retinal ganglion cells (RGCs) are the obligatory output neurons of the mammalian retina, with their axons constituting the principal components of the optic nerve. The optic nerve exits the posterior globe and projects to the brain, where RGC axons form synaptic connections with higher-order neurons. Along this pathway, RGC axons are influenced by two principal forces: intraocular pressure (IOP) acting at the intraocular and pre-lamina cribrosa regions, and intracranial pressure (ICP) acting posterior to the lamina cribrosa and along the optic nerve course [1, 2]. Idiopathic or secondary elevations in ICP or IOP can lead to vision-threatening pathologies, including spaceflight-associated neuro-ocular syndrome (SANS; ICP-related), idiopathic intracranial hypertension (IIH; ICP-related), and glaucoma (IOP-related) [3, 4]. The lamina cribrosa (LC), a critical anatomical structure, generates the translaminar cribriform pressure differential (TLCPD), which is determined by the interplay of IOP and ICP [5]. The pressure gradient between IOP and ICP is defined as the translaminar pressure difference (TLCPD or IOP-ICP).

TLCPD is hypothesized to play a pivotal role in optic nerve homeostasis and may contribute to the pathophysiology of certain optic neuropathies. For instance, SANS occurs under conditions of elevated ICP with normal IOP and reduced TLPD, resulting in optic disc edema and variable, often asymmetric, visual deficits [6]. Conversely, normal-tension glaucoma manifests with normal IOP but reduced ICP and elevated TLPD, leading to characteristic optic nerve cupping and visual field loss[6, 7]. Emerging evidence suggests that altered TLCPD induces structural deformation of the LC. Not surprisingly, several groups have proposed that a delicate equilibrium exists between IOP and ICP, where manipulation of one pressure may counteract dysregulation of the other[8, 9]. Unfortunately, direct measurement of human ICP remains technically challenging, necessitating invasive procedures such as lumbar puncture or intracranial instrumentation, which limits rigorous testing of these hypotheses, particularly in microgravity environments [10]. Although non-invasive ICP estimation methods exist, their clinical utility remains constrained[11]. Consequently, controlled experimental data on the effects of ICP on TLPD, optic nerve function, and biomechanics are scarce, largely due to difficulties in ICP measurement in humans and even model systems[5].

In the context of prolonged microgravity exposure, our research group has demonstrated that rats subjected to extended microgravity exhibit progressive deterioration of intraocular visual function [12]. However, the effects of microgravity on ICP and IOP regulatory mechanisms remain poorly characterized, further limiting mechanistic insights[13]. Our prior work established that experimental and chronic reductions in cerebrospinal fluid pressure (CSFP) induce optic neuropathy in rhesus macaques [14]. To date, no countermeasures have been implemented in spaceflight to mitigate SANS by rebalancing altered IOP and ICP in astronauts, underscoring the urgent need to elucidate pathophysiological mechanisms underlying visual impairment for effective ocular protection during space missions.

Elevated IOP-induced mechanical strain contributes to activation of retinal glial cells, including microglia and astrocytes. Early immune activation occurs prior to detectable morphological loss of RGCs [15, 16]. Activated microglia communicate with astrocytes, polarizing them into distinct phenotypes (A1 and A2), which exhibit neuroinflammatory or neuroprotective properties, respectively[17]. In our previous studies, a rat model of low ICP (LICP) induced by sustained CSFP reduction was well-validated, with selective early astrocytic reactivity observed in the retina of LICP rats[18]. Cytokines and neurotrophic factors released by glial cells may contribute to blood-retinal barrier (BRB) disruption and RGC injury [19]. Reactive glial cells can acquire antigen-presenting capabilities via upregulation of HLA-DR (a human MHC class II molecule), facilitating T-lymphocyte recruitment from peripheral circulation into the retina[20]. Intraocular dual-chamber pressure alterations have been shown to induce axonal and RGC degeneration, accompanied by CD4+ T-cell infiltration into the ganglion cell layer (GCL), indicative of adaptive immune involvement in visual dysfunction. Collectively, innate immune responses (microglial/astrocytic activation), BRB compromise, and adaptive immunity (T-cell infiltration) may represent three critical pathological steps driving neuroinflammation. However, the characteristics of microgravity-induced ocular inflammatory responses remain poorly understood.

Minocycline (Mino), a tetracycline antibiotic with BRB-penetrating properties, exhibits anti-inflammatory, anti-apoptotic, and antioxidant activities. It is widely employed to suppress neuroinflammation via inhibition of glial activation [21]. The therapeutic efficacy of Mino in ameliorating inflammatory cascades has been extensively documented in animal models of neurodegenerative disorders, including Alzheimer’s disease, diabetic retinopathy, retinal ischemia-reperfusion injury, and glaucoma[22–24]. Acetazolamide (AZE), an inhibitor of carbonic anhydrase (CA), targets the 15 CA isozymes ubiquitously expressed across mammalian cell types and tissues [25]. These enzymes catalyze the reversible hydration of CO2 to H2CO3, which subsequently dissociates into H+ and bicarbonate ions. Initially introduced in the 1950s as a diuretic, AZE was later repurposed for managing elevated ICP [26]. It remains a first-line therapy for select ICP-related pathologies, such as idiopathic intracranial hypertension [27], irrespective of its precise mechanism in ICP modulation. Previous studies have validated AZE’s efficacy in reducing ICP in freely moving male rats[28] [29].

This study aims to investigate the protective effects of different interventions on visual function in simulated microgravity rats, focusing on the impact of AZE in modulating the intraocular mechanical environment and the mechano-neuroimmune crosstalk effects on visual parameters following Mino-mediated neuroimmune suppression.

## Materials and Methods

### 1. Ethical Approval and Grouping Protocol

The experimental protocol was approved by the Animal Care and Use Committee of Capital Medical University (Approval No. AEEI-2024-030) and conducted in strict compliance with the Association for Research in Vision and Ophthalmology (ARVO) Statement for the Use of Animals in Ophthalmic and Vision Research. Forty-eight male Sprague-Dawley rats (weight range: 250–300 g), obtained from Beijing Vital River Laboratory Animal Technology Co., Ltd., underwent stratified randomization into two primary groups: a ground-based control group (CTRL, *n* = 12) and a simulated microgravity group (*n* = 36) induced via hindlimb suspension (HLS). The HLS group was further subdivided into three intervention-based subgroups— phosphate-buffered saline (PBS), acetazolamide (AZE), and minocycline (MINO)—each stratified by exposure duration (2 weeks or 4 weeks), yielding six experimental subgroups: PBS-2W (*n* = 6), PBS-4W (*n* = 6), AZE-2W (*n* = 6), AZE-4W (*n* = 6), MINO-2W (*n* = 6), and MINO-4W (*n* = 6).

### 2. Establishment of Simulated Microgravity Model

The hindlimb suspension (HLS) model, a well-established method for simulating microgravity, was implemented as previously described [30]. Following a 7-day acclimatization period, the rats underwent tail preparation: the tail surface was cleaned and coated with a mixture of benzoin tincture and rosin ethanol solution to enhance adhesive tape adherence and prevent tissue injury. After complete drying, medical-grade adhesive tape (20 cm in length) was affixed to the tail, enabling suspension at a 30° head-down tilt within customized cages. The forelimbs remained freely mobile to allow access to food and water, while hindlimb contact with the cage floor was systematically restricted. Throughout the experiment, environmental parameters were strictly controlled (temperature: 23 ± 1°C; relative humidity: 45–55%; light-dark cycle: 12 h/12 h).

### 3. Drug Preparation

Acetazolamide (HY-B0782, Acetazolamide, Monmouth Junction, NJ, USA) was administered to investigate its protective effects on visual function in simulated microgravity rats by modulating intraocular pressure (IOP) and intracranial pressure (ICP). The HLS group received daily intragastric gavage at a fixed time (09:00–10:00) with a dosage of 40 mg/100g. The working solution was prepared as follows: 100 μL of 25.0 mg/mL dimethyl sulfoxide (DMSO) stock solution was mixed with 400 μL of 40% PEG300, followed by the addition of 50 μL of 5% Tween-80. Subsequently, 450 μL of 45% physiological saline (prepared by dissolving 0.9 g sodium chloride in deionized water [ddH₂O], adjusted to 100 mL) was added to achieve a final volume of 1 mL. This formulation yielded a clear solution with a concentration ≥2.5 mg/mL (11.25 mM; solubility ≥2.5 mg/mL, saturation unspecified).

Minocycline (CAS No. 13614-98-7, R013129, Rhawn, Shanghai, China) was utilized to assess its neuroprotective effects in rats with low intracranial pressure (LICP). The treatment regimen involved an initial intraperitoneal injection of 5 mg/100g, followed by subsequent injections every 48 hours at a reduced dose of 2.5 mg/100g [30]. For solution preparation, minocycline was dissolved in phosphate-buffered saline (PBS) via ultrasonic-assisted dissolution until a clear solution with a solubility of 7.69 mg/mL (15.57 mM) was obtained. This protocol ensured complete dissolution of minocycline in PBS, yielding a stable solution for experimental use.

### 4. Intraocular Pressure (IOP) Measurement

Intraocular pressure (IOP) was measured using a rebound tonometer (TonoLab, Icare, Helsinki, Finland) in "rat" mode. Prior to measurement, rats were anesthetized with 3.5% isoflurane until loss of self-righting reflex (typically achieved within 4–5 minutes). IOP measurements were conducted daily between 09:00 and 11:00 AM. The tonometer probe was positioned perpendicular to the corneal surface, and five consecutive readings were obtained per eye, with the average value calculated. To ensure accuracy, this process was repeated three times for each eye, and the mean of these three averaged values was recorded as the final IOP

### 5. Intracranial Pressure (ICP) Measurement

Intracranial pressure (ICP) monitoring was performed under 3% isoflurane anesthesia, with depth of anesthesia confirmed by loss of self-righting reflex and absence of hindlimb withdrawal response to painful stimuli. Following hair removal and disinfection of the surgical site, a midline scalp incision was made to expose the skull by dissecting fascia and muscle layers. Using stereotaxic guidance, the Bregma landmark was identified, and a precise drilling site was determined (1.5 mm posterior to Bregma, 0.8 mm lateral), followed by a 4 mm deep craniotomy[31]. A 1 mm diameter dental drill (Dremel, Racine, WI, USA) was used to gradually penetrate the bone. Upon confirmation of dura mater rupture, a 3 mm pressure transducer was advanced 1 mm into the cerebral parenchyma. The transducer was connected via a dedicated amplifier (Bridge Amp, AD Instruments, Australia) to a biosignal acquisition system (PowerLab 16/35, AD Instruments, Australia) for continuous ICP monitoring at a sampling rate of 1,000 Hz (units: mmHg). After signal stabilization, three consecutive measurement cycles were conducted to obtain the mean ICP value. Postoperative care included saline irrigation, layered suturing (muscle, fascia), and skin closure under aseptic conditions. Rats were allowed to recover on a thermostatically controlled heating pad for approximately 20 minutes until full consciousness was regained.

### 6. Translaminar Cribriform Pressure Difference (TLCPD) Measurement

The translaminar cribriform pressure difference (TLCPD) was defined as the differential pressure between intraocular pressure (IOP) and intracranial pressure (ICP). IOP and ICP measurements were conducted using the standardized protocols detailed in Sections 3 and 4, respectively. TLCPD values (expressed in millimeters of mercury, mmHg) were calculated by subtracting the mean ICP value from the corresponding mean IOP value for each eye. Subsequent statistical analyses were performed to evaluate TLCPD variations across experimental groups and time points.

### 7. Electrophysiological Assessment (ERG)

Retinal electrophysiological evaluation was performed using an Espion Visual Electrophysiology System (Diagnosys LLC, USA). Following a minimum of 6 hours of dark adaptation, pupillary dilation was achieved by topical application of a compound tropicamide solution (Santen Pharmaceutical Co., Ltd., Japan). During recording, rats were positioned on a thermostatically controlled heating pad to maintain physiological body temperature. Electrode configuration included a gold-ring corneal contact electrode (interface lubricated with carbomer gel; Bausch & Lomb, Jinan, China), with subcutaneous reference and ground electrodes inserted at the cheek and tail, respectively. Dark-adapted full-field flash stimuli (3.0 cd·s/m²) were delivered to elicit characteristic waveforms, with a focus on analyzing the amplitudes of the a-wave, b-wave, and photopic negative response (PhNR). The PhNR component, emerging post-b-wave, predominantly reflects retinal ganglion cell (RGC) activity. Bilateral waveform amplitudes were systematically recorded for subsequent quantitative analysi

### 8. Optomotor Response (OMR)

Visual function was assessed using a virtual optomotor tracking system (OptoMotry, Cerebral Mechanics Inc., Lethbridge, Canada). Rats were positioned on a central platform within a testing chamber surrounded by four computer monitors displaying rotating virtual spatial frequency gratings. A ceiling-mounted camera within the chamber monitored rodent behavior. A psychophysical staircase paradigm was employed, wherein spatial frequency was randomly adjusted until the threshold stimulus eliciting a visual motion response under optimal contrast was determined. To independently evaluate monocular visual function, gratings were rotated in clockwise or counterclockwise directions to selectively stimulate the contralateral eye. Mean spatial frequency thresholds for each eye were recorded and analyzed.

### 9. Optical Coherence Tomography (OCT)

Retinal structural evaluation was performed using the Micron III OCT system (Phoenix Research Labs, Pleasanton, CA, USA). Under standardized anesthesia, pupillary dilation was achieved with 0.2 mg/mL phenylephrine, and corneal hydration was maintained using 1.5% hydroxyethyl cellulose solution. Horizontal OCT scans centered on the optic nerve head (ONH) were acquired via an integrated fundus camera, with scan orientation marked by reference lines. To enhance image quality, 30 consecutive scans were averaged to minimize projection artifacts. Quantitative morphometric analysis was conducted using ImageJ software (National Institutes of Health, Bethesda, MD, USA). Layer-specific measurements of the retina were performed at standardized locations (200 µm nasal and temporal to the ONH), including total retinal thickness, NFL-GCL (nerve fiber layer-ganglion cell layer) complex thickness, inner nuclear layer (INL), and outer nuclear layer (ONL).

### 10. Immunofluorescence

Following enucleation, rat eyes were fixed overnight in 4% paraformaldehyde (PFA) at 4°C. For retinal flat-mount preparation, the cornea, lens, and vitreous body were carefully removed, and the isolated retinas were immersed in 0.5% Triton X-100 for 4 hours. Subsequently, retinas were blocked with 1% bovine serum albumin (BSA, A8010, Solarbio, Beijing, China) for 2 hours and incubated overnight at 4°C with the following primary antibodies: anti-RNA-binding protein with multiple splicing (RBPMS, 1:500, GTX118619, GeneTex, Irvine, CA, USA), anti-ionized calcium-binding adapter molecule 1 (Iba-1, 1:500, 019-19741, Fujifilm Wako Pure Chemical Corporation, Japan), and anti-glial fibrillary acidic protein (GFAP, 1:500, #3670, Cell Signaling Technology, Danvers, MA, USA). Iba-1, a microglia-specific calcium-binding protein, served as a marker for microglial activation. The following day, retinas were incubated with fluorescent secondary antibodies for 1 hour at room temperature under dark conditions, followed by nuclear counterstaining with DAPI (ab104139, Abcam, Cambridge, UK). Non-overlapping images of the entire retina were captured using a fluorescence microscope (Leica Microsystems, Wetzlar, Germany). RBPMS-stained retinal ganglion cells (RGCs) and Iba-1-positive microglia were quantified using ImageJ software (v1.53, National Institutes of Health, Bethesda, MD, USA) to systematically assess neural and immune alterations under experimental conditions.

### 11. Microglial Morphological Assessment

Microglial morphology was quantitatively analyzed using Sholl analysis to determine the number of dendritic intersections at varying radial distances [31, 32]. In FIJI (ImageJ), a straight line was drawn from the centroid of Iba-1-positive cells to the distal tip of the longest dendrite. Green fluorescence-labeled microglia were isolated via the Image > Color > Split Channels function. Morphological enhancement was achieved by applying Process > Filters > Unsharp Mask and Noise > Despeckle to clarify dendritic structures. Subsequently, images were converted to binary format using Image > Adjust > Threshold to enable fluorescence quantification. Sholl analysis was executed via Neuroanatomy Shortcuts > Sholl Analysis > Legacy: Sholl Analysis (From image), yielding parameters including mean dendritic length, maximum dendritic extension, and branchpoint density.

### 12. Statistical Analysis

Statistical analyses were performed using SPSS 27.0 (IBM Corp., Armonk, NY, USA), with graphs generated in GraphPad Prism 9.0 (GraphPad Software, San Diego, CA, USA). Data normality was verified via Shapiro-Wilk tests. Longitudinal effects of simulated microgravity were assessed across time points (2W, 4W, 8W) using one-way ANOVA. For ANOVA-significant effects, post hoc Dunnett’s tests compared experimental subgroups to the PBS control, while inter-time-point differences were evaluated with independent-sample t-tests. Non-normally distributed data were analyzed via Kruskal-Wallis tests, followed by Dunn’s post hoc comparisons. All post hoc analyses employed Bonferroni correction. Data are expressed as mean ± standard deviation (SD), with statistical significance set at P < 0.05

## Result

### 1. Effects of Interventions on Ocular Dual-Compartment Pressures

In this study, we first assessed dual-compartment pressure dynamics across experimental groups. Biomechanical alterations were most pronounced in rats under simulated microgravity. For intraocular pressure (IOP), both PBS and MINO groups exhibited significant elevations at 2W and 4W (p < 0.05), with greater increases at 4W compared to 2W (p < 0.05), indicating progressive microgravity-induced ocular hypertension (figure1.A). In contrast, AZE-treated rats showed statistically significant IOP reduction at 2W. By 4W, despite sustained IOP elevation in other groups, AZE maintained IOP levels comparable to baseline, demonstrating significant differences versus PBS and MINO groups (p < 0.05).For intracranial pressure (ICP), PBS-treated microgravity rats displayed slight non-significant increases at 2W and 4W, with IOP rising more sharply than ICP. (figure1.B) Comparative analysis revealed significant differences between 2W and 4W dual-pressure profiles (p < 0.05), suggesting ocular biomechanical remodeling occurs within short-term microgravity exposure. AZE significantly reduced both IOP and ICP from treatment initiation to 4W (p < 0.05), though ICP reduction was less pronounced than IOP. MINO group pressures remained comparable to PBS (P > 0.05), differing significantly only from ground controls and PBS groups (p < 0.05).Translaminar cribriform pressure difference (TLCPD) dynamics were prioritized. In controls, ICP marginally exceeded IOP, establishing the baseline pressure gradient direction. (figure1.C) Short-term microgravity exposure reversed this gradient in PBS rats due to disproportional IOP elevation versus ICP. AZE reduced TLCPD magnitude by preferentially lowering IOP, though the mechanical gradient direction remained unchanged. By 4W, AZE exhibited significantly reduced absolute TLCPD compared to MINO and PBS groups (p < 0.05)

**Figure 1.**
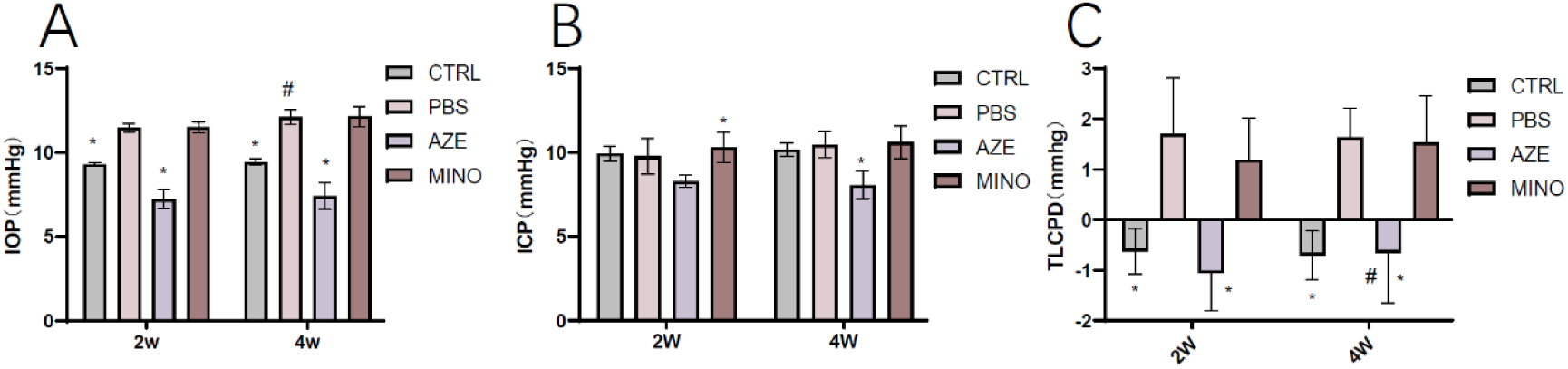
Intervention effects on intraocular pressure, intracranial pressure, and translaminar cribriform pressure difference in simulated microgravity rats. Intraocular pressure (IOP) values (n = 6). (B) Intracranial pressure (ICP) values (n = 6). (C) Translaminar cribriform pressure difference (TLCPD) in simulated microgravity rats (n = 6). SW indicates simulated microgravity. **P < 0.01, #Significant difference between 2-week and 4-week timepoints within the same treatment group (P < 0.05)

### 2. Functional Visual Protection: ERG and OMR

To evaluate the protective effects of interventions on retinal function under simulated microgravity, electroretinogram (ERG) and optomotor response (OMR) spatial frequency analyses were performed in rats exposed to simulated weightlessness for 2 and 4 weeks (Figures 3A and 3B). During OMR testing, all hindlimb-suspended rats initially exhibited locomotor hypoactivity, compromising behavioral accuracy. To mitigate this, the acclimatization period was extended from 15 to 30 minutes, yielding reliable contrast sensitivity data. Results demonstrated no significant behavioral differences in the 2-week microgravity group (P > 0.05). However, by 4 weeks, the PBS group showed statistically significant contrast sensitivity reduction (P < 0.05).

The AZE group exhibited marked sensitivity decline compared to 2-week timepoints (P < 0.05), while the MINO group maintained baseline performance. (figure2.D) In ERG assessments, the a-wave (photoreceptor function) displayed non-significant amplitude reductions in the PBS group at 2W and 4W compared to controls, potentially due to limited exposure duration. AZE-treated rats showed partial a-wave recovery, though not reaching control levels, likely attributable to altered intraocular biomechanics. The MINO group, benefiting from neuroimmune protection, maintained higher a-wave amplitudes than PBS at both timepoints, albeit without statistical significance.The b-wave (bipolar cell–retinal ganglion cell transmission) declined significantly in controls and PBS groups (P < 0.05), with greater reductions at 4W versus 2W. AZE conferred minimal protection at 2W, but progressive efficacy emerged by 4W as injury severity increased, though differences from PBS remained non-significant. In contrast, MINO significantly preserved b-wave amplitudes versus PBS (P < 0.05).The photopic negative response (PhNR; retinal ganglion cell action potentials) paralleled b-wave trends. PhNR amplitudes decreased significantly at 2W and 4W across groups (P < 0.05). Notably, MINO-treated rats maintained PhNR amplitudes comparable to ground controls, demonstrating significant protection versus PBS and AZE groups (P < 0.05), thereby underscoring MINO’s efficacy in preserving visual function under simulated microgravity. (figure2.A-C)

**Figure 2.**
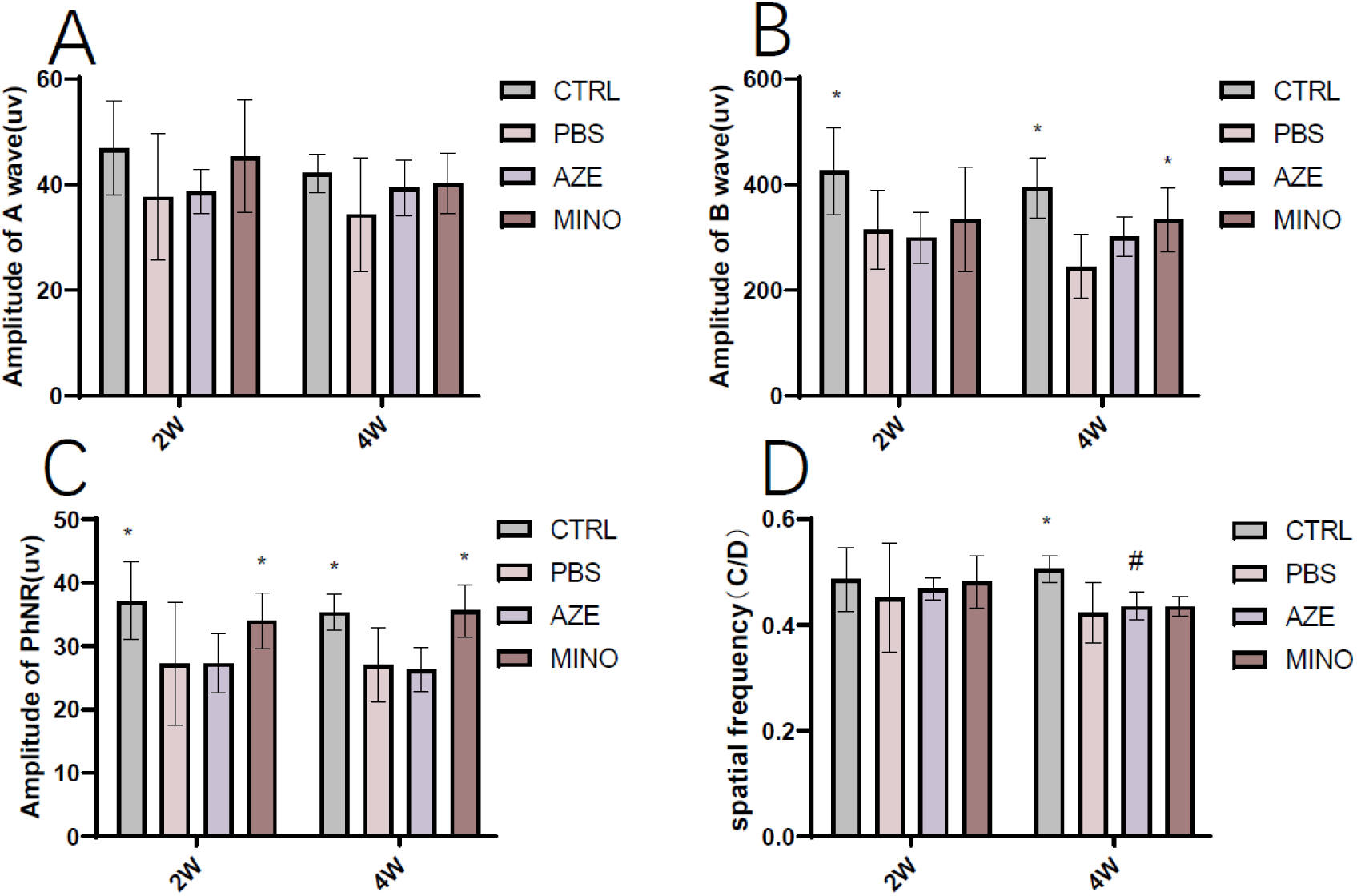
Visual Function Outcomes in Simulated Microgravity Rats. (A-C) Representative waveforms of a-wave, b-wave, and photopic negative response (PhNR) from electroretinogram (ERG) recordings. (D) Spatial frequency contrast sensitivity (cycles/degree, C/D) assessed by optomotor response (OMR) testing in dark-adapted simulated microgravity rats (n = 6). *P < 0.05 vs. control group at the same timepoint; #P < 0.05 vs. control group at the same timepoint.

### 3. In Vivo Retinal OCT Measurements

This study employed optical coherence tomography (OCT) to quantify retinal layer thickness, revealing spatiotemporally heterogeneous structural damage in rat retinas under simulated microgravity. (figure3.A-D) Total retinal thickness showed no significant differences between 2-week (2W) and 4-week (4W) microgravity exposure (P > 0.05), suggesting short-term exposure did not induce global structural alterations. Consequently, sublayer-specific analyses were prioritized.Nerve Fiber Layer–Ganglion Cell Layer (NFL-GCL) Complex: The PBS group exhibited significant NFL-GCL thickening at 4W compared to 2W (P < 0.05), whereas AZE and MINO groups maintained thickness comparable to controls. This implies that biomechanical modulation (AZE) and neuroimmune suppression (MINO) may mitigate microgravity-driven axonal remodeling in the outermost retinal layer.Inner Nuclear Layer (INL): No significant INL changes were observed at 2W across groups. By 4W, the PBS group showed increased INL thickness compared to intervention groups (P < 0.05), though within-group differences (2W vs. 4W) were non-significant. This suggests potential metabolic dysregulation in bipolar/Müller cells under prolonged microgravity.Outer Nuclear Layer (ONL): At 2W, both MINO and AZE groups displayed ONL thinning versus controls—a divergence from other layers. By 4W, this trend attenuated, with AZE showing significant ONL preservation compared to PBS (P < 0.05), indicating neuroinflammatory suppression (MINO) may substantially protect against photoreceptor degeneration.

**Figure 3.**
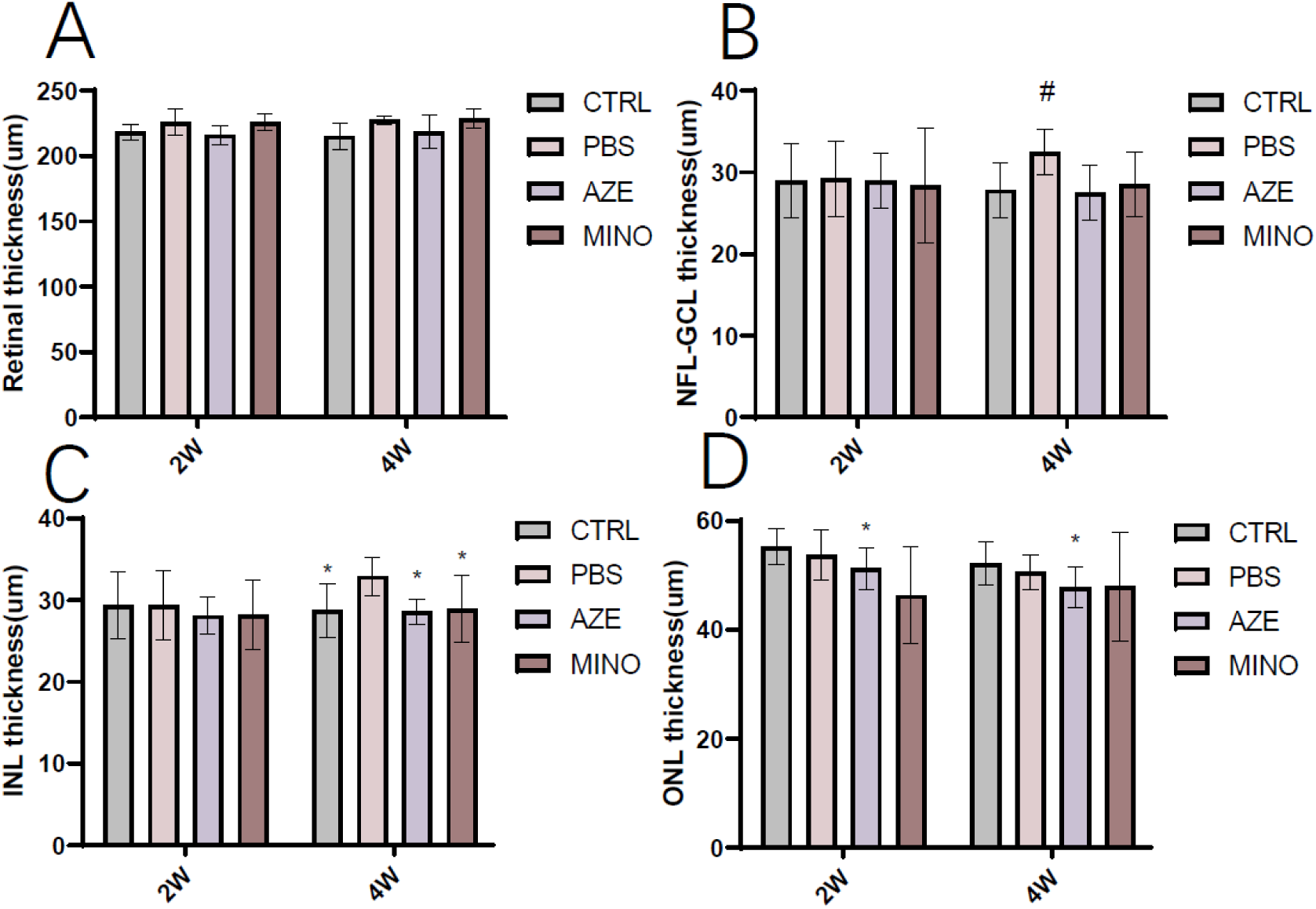
Retinal thickness measurements via optical coherence tomography (OCT) in simulated microgravity rats. (A) Total retinal thickness (µm). (B) Nerve fiber layer-ganglion cell layer complex (NFL-GCL) thickness (µm). (C) Inner nuclear layer (INL) thickness (µm). (D) Outer nuclear layer (ONL) thickness (µm) (n = 6). *P < 0.05 vs. control group at the same timepoint; #P < 0.05 vs. control group at the same timepoint. SW indicates simulated microgravity group

### 4. Retinal Fundus Imaging

Fundus imaging analyses revealed progressive structural alterations in rat retinas under simulated microgravity (Figure 4A-H). High-resolution fundus photography demonstrated marked optic disc hyperemia across all microgravity groups versus controls. In the PBS group, retinal vasculature remained structurally intact during 2-week exposure, with no significant choroidal vascular changes. AZE-treated rats showed comparable vascular morphology to other groups but exhibited reduced choroidal congestion, likely attributable to AZE-mediated pressure reduction. The MINO group remained comparable to controls. (figure4.C.G) By 4 weeks, PBS rats displayed pronounced choroidal vascular engorgement with mild vessel thickening (no tortuosity), suggesting pressure-driven vascular distension during short-term exposure. In AZE rats, pallor severity decreased versus 2W, accompanied by partial choroidal congestion alleviation; however, optic disc hyperemia persisted without significant improvement. The MINO group mirrored PBS outcomes. These findings suggest MINO may preserve choroid-retinal hemodynamic homeostasis via neuroinflammatory suppression, whereas AZE’s biomechanical modulation—while partially normalizing dual-compartment pressures—offers limited protection to photoreceptor microarchitecture. (figure4.D.H)

**Figure 4.**
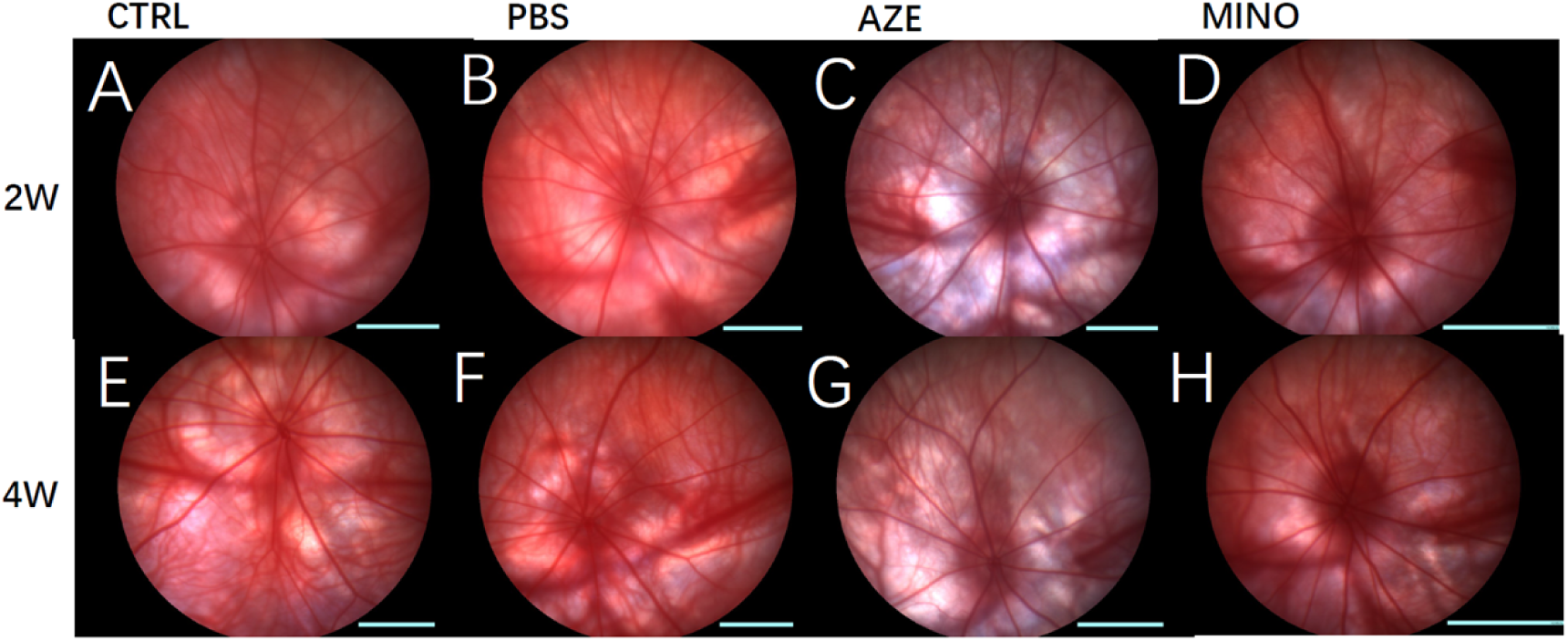
Longitudinal fundus imaging in simulated microgravity rats. Representative in vivo fundus photographs of (A-D) 2-week and (E-H) 4-week exposure groups

### 5. Retinal Ganglion Cell (RGC) Quantification

To evaluate the effects of microglial inhibition on neuroinflammation and RGC function, MINO was administered via intraperitoneal injection. RGC density changes across exposure durations were systematically quantified using retinal flat-mounts combined with RBPMS immunofluorescence staining (Figure 5A-H). Short-term microgravity exposure (2W and 4W) induced no significant RGC loss in the PBS group compared to ground controls (CTRL). However, the AZE group exhibited significant RGC reduction at 2W versus CTRL and PBS groups (P < 0.05). By 4W, AZE-treated rats showed partial RGC recovery, reaching levels comparable to CTRL and PBS but retaining statistical significance (P < 0.05). Notably, the MINO group displayed marginally higher RGC counts than CTRL, though high interindividual variability (reflected in large SD) suggested heterogeneity in degenerative progression, potentially linked to the pathological sequence where axonal injury precedes somatic loss. These findings aligned with progressive PhNR amplitude attenuation in ERG, indicating that prolonged microgravity exposure drives irreversible RGC loss via chronic mechanical-inflammatory stress. Both interventions (AZE, MINO) yielded numerically higher RGC counts than PBS, but differences lacked statistical significance (P > 0.05). (Figure 5.I)

**Figure 5.**
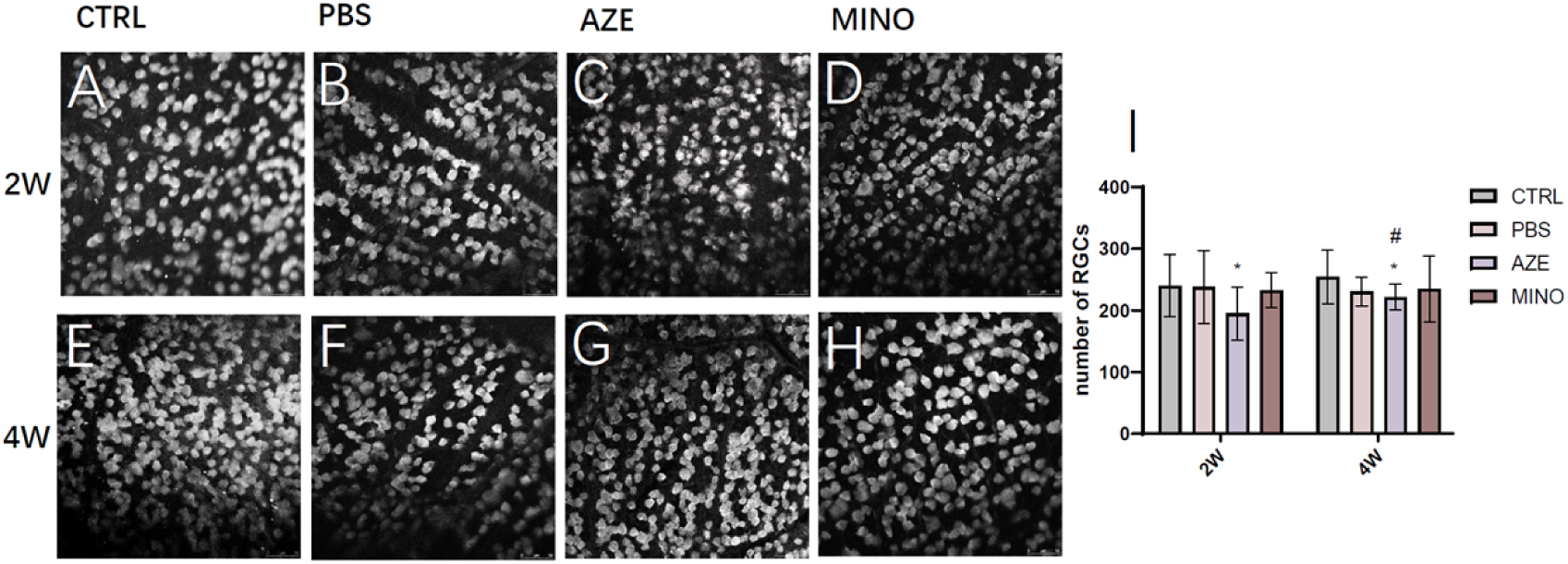
Immunofluorescence analysis of retinal changes under simulated microgravity. Representative retinal flat-mount images showing RBPMS immunostaining (gray) of retinal ganglion cells (RGCs) (A-D) 2-week exposure groups (E-H) 4-week exposure groups Experimental groups: ground control, PBS-treated, AZE-treated, MINO-treated; n = 3 eyes per group/timepoint).(I) Quantitative analysis of RGC density (cells/mm²). RGC, retinal ganglion cell; SW, simulated microgravity group. *P < 0.05 vs. control group at the same timepoint; #P < 0.05 vs. control group at the same timepoint.

### 6. Astrocytic Activation

GFAP immunofluorescence staining demonstrated time-dependent activation of astrocytes under simulated microgravity, with MINO intervention significantly attenuating this pathological process, while AZE showed no notable protective effects comparable to the PBS group. (Figure 6A-H)At the 2-week exposure timepoint, PBS-treated rats exhibited early reactive gliosis characterized by mild astrocytic soma hypertrophy, subtle thickening of cellular processes, and modest upregulation of GFAP expression. In contrast, MINO-treated rats maintained astrocytic morphology and GFAP expression levels similar to ground controls, indicating effective suppression of initial glial activation. By 4 weeks, PBS rats displayed pronounced astrocytic activation, marked by increased branching complexity, significant process thickening, and robust GFAP overexpression, suggesting peak glial reactivity that could exacerbate neuroinflammatory cascades and compromise blood-retinal barrier integrity. Remarkably, MINO-treated rats retained near-normal astrocytic morphology with only mild GFAP elevation, demonstrating its capacity to inhibit pathological hyperactivation through anti-inflammatory mechanisms. Furthermore, MINO modulated astrocytic phenotypes by suppressing pro-inflammatory transitions, thereby preserving the functional integrity of the glia-RGC interaction network and creating a protective microenvironment conducive to RGC survival. These findings underscore the divergent efficacy of interventions: MINO’s neuroimmune suppression effectively disrupted chronic inflammatory stress, whereas AZE’s biomechanical modulation failed to mitigate glia-driven retinal damage.。

**Figure 6.**
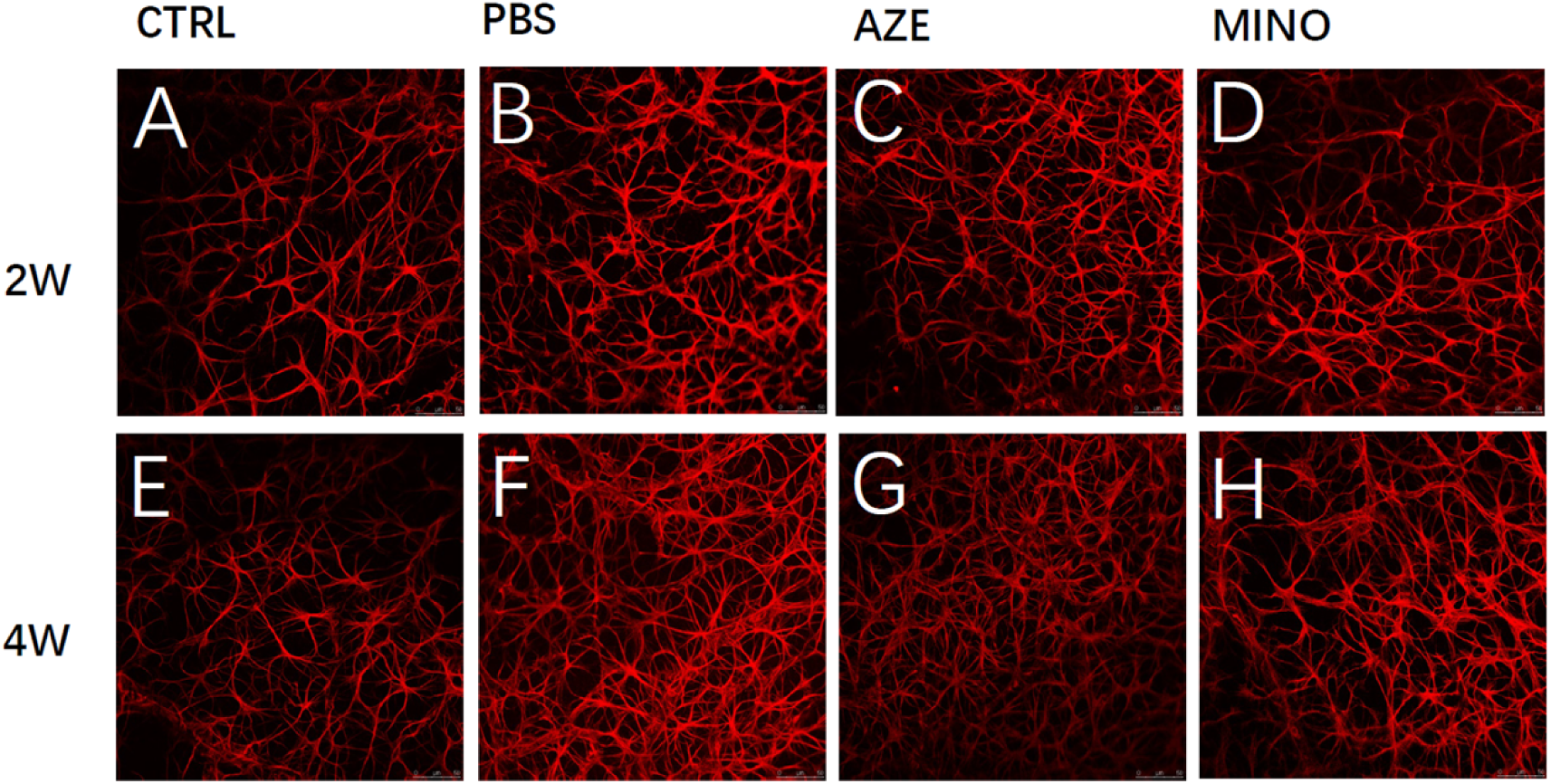
Retinal astrocyte activation and morphological changes under simulated microgravity. Retinal flat-mount images showing glial fibrillary acidic protein (GFAP) immunostaining (red) of astrocytes:**A-D**: 2-week exposure groups**E-H**: 4-week exposure groups.Experimental groups: ground control, PBS-treated, AZE-treated, MINO-treated; n = 3 eyes per group/timepoint).

### 7. Microglial Activation and Sholl Analysis

To investigate the impact of microglial inhibition on neuroinflammation and RGC function, MINO was administered via intraperitoneal injection, with microglial morphological dynamics assessed through Iba-1 immunofluorescence. (Figure 7A-H) Compared to ground controls, the PBS group exhibited a slight increase in Iba-1⁺ microglia at 2 weeks, predominantly retaining ramified morphologies suggestive of early activation. By 4 weeks, PBS rats showed a significant surge in microglial density, accompanied by characteristic morphological shifts: soma hypertrophy, process retraction, and shortened branches. In contrast, the MINO group displayed markedly fewer Iba-1⁺ cells and reduced activation severity. Sholl and skeleton analyses quantified these morphological alterations. Sholl analysis revealed that PBS rats at 2 weeks had significantly fewer dendritic intersections (P < 0.05) and shorter average branch lengths compared to controls, reflecting a transition from resting to activated states. (Figure 8A-H) By 4 weeks, microglial activation intensified, with further soma enlargement, reduced branching complexity, and a subset adopting dysfunctional “amoeboid-like” morphologies. Skeleton analysis corroborated these findings, demonstrating significant reductions in branch number and length at both 2 and 4 weeks in PBS rats, indicative of microglial functional impairment under prolonged microgravity. Conversely, MINO treatment significantly restored branch length and complexity at both timepoints (P < 0.05). These collective results highlight microglial activation as a pivotal driver of simulated microgravity-induced retinal degeneration, with MINO effectively suppressing this pathogenic activation to preserve retinal integrity

**Figure 7.**
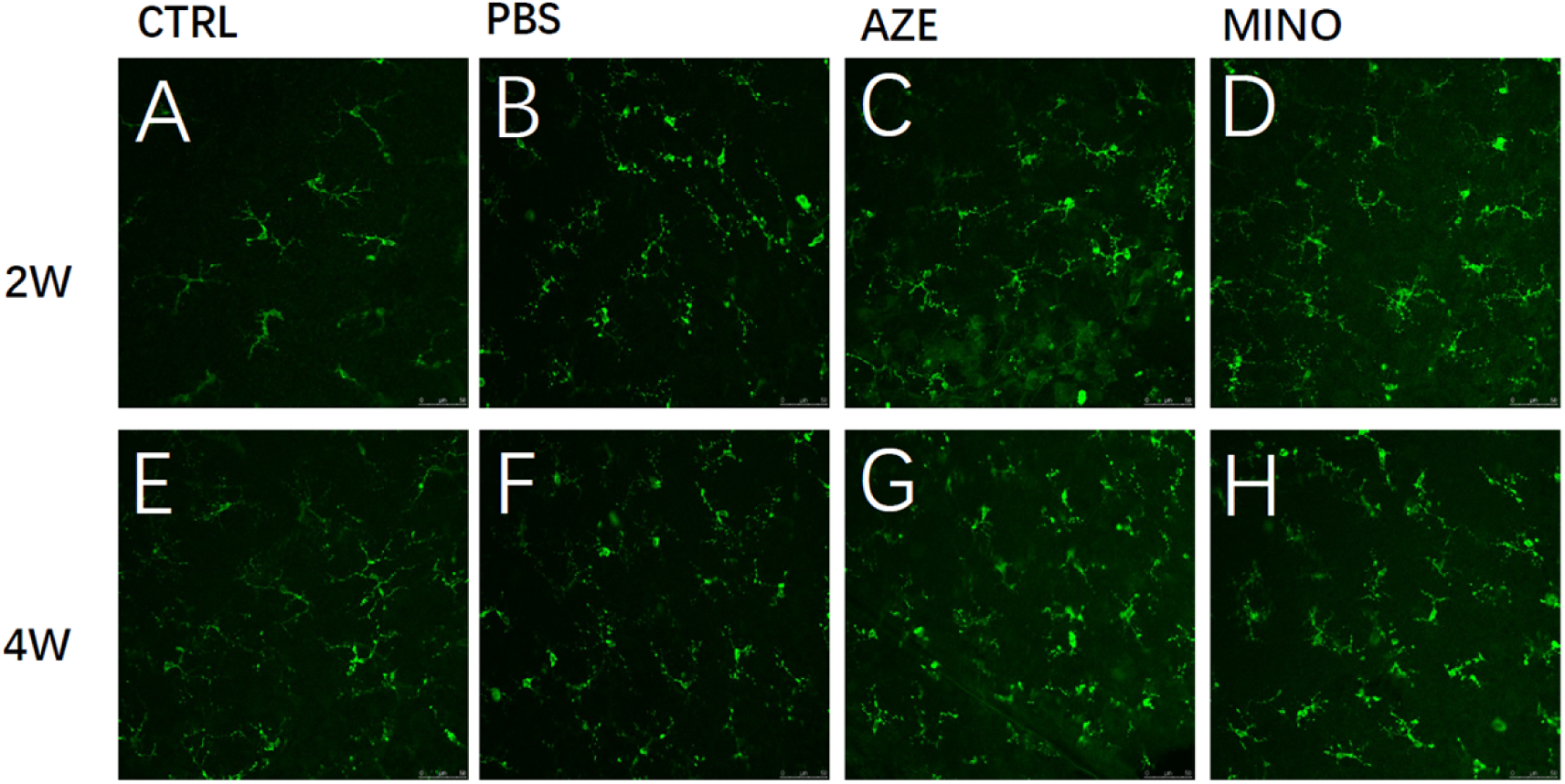
Microglial activation and morphological remodeling in simulated microgravity rat retinas. **(A-H)** Retinal flat-mount images showing ionized calcium-binding adapter molecule 1 (IBA-1) immunostaining (green) of microglia:**A-D**: 2-week exposure groups.**E-H**: 4-week exposure groups .Experimental groups: ground control, PBS-treated, AZE-treated, MINO-treated; n = 3 eyes per group/timepoint).

**Figure 8.**
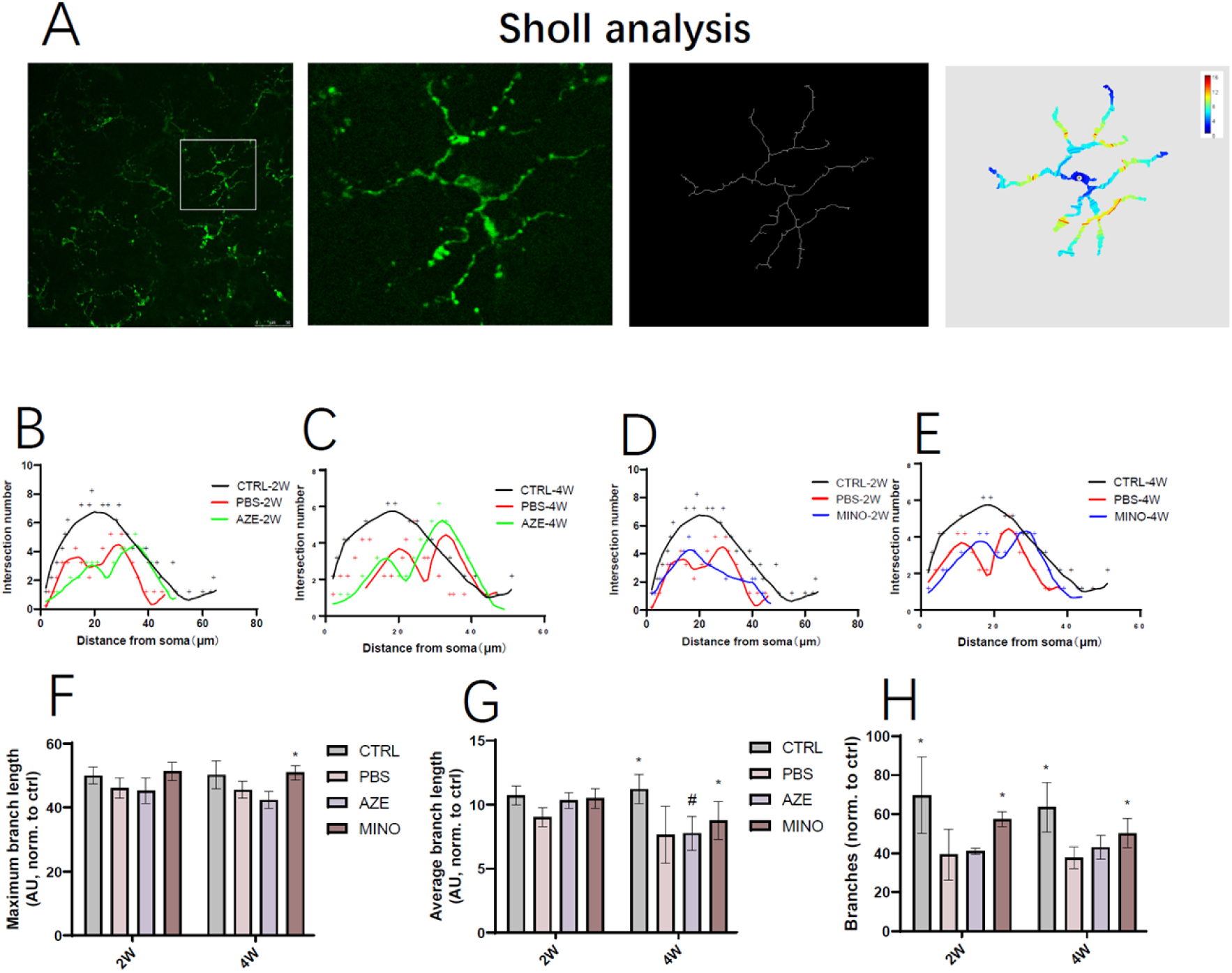
Sholl analysis of microglial morphological complexity in simulated microgravity rat retinas. (A) Sholl analysis workflow: Representative Iba-1⁺ microglia (green) were binarized and processed for concentric circle intersection analysis at 2-week and 4-week exposure timepoints (n = 3 eyes per group/timepoint). (B-E) Quantification of Sholl intersections across experimental groups. (F) Maximum branch length, (G) mean branch length, and (H) total branch number. SW, simulated microgravity group. *P < 0.05 vs. control group at the same timepoint; #P < 0.05 vs. control group at the same timepoint.

## Discussion

The lamina cribrosa (LC), a porous connective tissue network within the optic nerve head, provides mechanical support for retinal ganglion cell (RGC) axons and serves as a pressure-sensitive structure [32]. The LC exhibits high sensitivity to the synergistic effects of intraocular pressure (IOP) and intracranial pressure (ICP). Mechanical deformation of the LC triggers pathological cascades, including axoplasmic transport disruption of neurotrophic factors, tissue hypoxia, and glial activation, ultimately leading to neuronal death [33]. This mechanism shares commonality across glaucoma, idiopathic intracranial hypertension (IIH), and spaceflight-associated neuro-ocular syndrome (SANS). For instance, elevated ICP in IIH patients impedes neurotrophic signaling across the LC, while optic nerve damage in glaucoma and irreversible visual dysfunction in SANS are closely linked to abnormal translaminar pressure gradients (IOP-ICP differentials) [34] Microgravity-induced cephalad fluid redistribution drives multisystem adaptations, including cardiovascular dysregulation and intracranial hemodynamic abnormalities [35]. Our study revealed that simulated microgravity provoked a mild elevation in intraocular pressure (IOP) at 2 weeks, with further amplification observed at 4 weeks, consistent with prior reports attributing acute IOP spikes to choroidal congestion during short-term microgravity exposure [36]. Prolonged exposure likely triggers chronic pathological IOP elevation through increased episcleral venous pressure and augmented aqueous humor outflow resistance. Notably, intracranial pressure (ICP) remained stable under short-term (2- and 4-week) simulated microgravity in our model, suggesting compensatory cerebrospinal fluid (CSF) redistribution may mitigate intracranial pressure fluctuations. However, in-flight astronaut data demonstrate more complex pressure dynamics: Lee et al. [37] reported post-mission lumbar CSF pressures of 28–28.5 cm H₂O (normal range: 5–15 cm H₂O) in two spaceflight-associated neuro-ocular syndrome (SANS) patients, which persisted at 21–22 cm H₂O even after 66 days of recovery—substantially exceeding ground-based predictions, potentially due to impaired jugular venous return and interspecies differences in CSF compensatory mechanisms [38]. This pressure disequilibrium exacerbates optic neuropathy via dual mechanisms: (i) Elevated ICP induces optic nerve sheath distension and axonal myelin edema, synergizing with oligodendrocyte trophic insufficiency to drive demyelination; (ii) Altered IOP-ICP gradients directly strain the lamina cribrosa (LC), redistributing mechanical stress to disrupt axonal microarchitecture. Microgravity-associated intracranial vascular remodeling may critically modulate ICP regulation. Long-duration spaceflight induces gray/white matter volumetric anomalies and CSF space expansion[39], while Kergoat’s retinal ischemia hypothesis [40]posits that even transient microgravity disrupts retinal hemodynamics, synergizing with translaminar pressure gradients to accelerate SANS progression. Although our rat model exhibited limited ICP elevation, the interplay between IOP dynamics and residual ICP shifts significantly modulated LC biomechanics. Chronic mechanical stress may activate Piezo1 channels in astrocytes, triggering NF-κB-mediated inflammatory cascades that compromise blood-retinal barrier integrity and visual function[41] Despite the absence of a human-like collagenous LC structure in rodent optic nerve heads, astrocyte-derived cribriform analogs partially mediate IOP/ICP mechanotransduction[42]. This anatomical divergence requires nuanced interpretation: On one hand, the exclusive reliance on central retinal artery perfusion (rather than choroidal circulation) in rodents[43] renders them a unique model for studying independent IOP/ICP regulatory mechanisms. Conversely, simulated microgravity induced progressive translaminar cribriform pressure difference (TLCPD) shifts—significant anterior displacement at 2 weeks, intensifying by 4 weeks—without corresponding functional recovery (e.g., unaltered ERG waveforms). Canine models confirm linear correlations between optic nerve surface deformation and IOP-ICP gradients[44], yet real-time microstructural responses of LC components (e.g., collagen bundle thickness, pore geometry) to combinatorial pressure stimuli remain uncharacterized. Chronic pressure dysregulation may induce LC remodeling (e.g., altered collagen crosslinking density), obscuring distinctions between acute biomechanical insults and chronic adaptive changes.

As theorized earlier, perturbations to pressure homeostasis may manifest in three potential scenarios: (1) Directional TLCPD reversal with stable magnitude; (2) Magnitude alteration without directional shift; (3) Baseline pressure modifications (IOP/ICP) without TLCPD alteration[33] [34]. Our study primarily observed Scenario 1 (increased TLCPD magnitude), whereas AZE intervention induced Scenario 2 (direction reversal with elevated magnitude). Complementary studies in hypo-ICP and hyper-IOP models demonstrate that TLCPD reduction below critical thresholds disrupts retinal ganglion cell (RGC) axonal transport and mitochondrial function[18, 45], further implicating biomechanical-neuroimmune crosstalk in microgravity-associated intraocular dysfunction.

This study systematically investigated dynamic changes in intraocular pressure (IOP), visual function, and retinal structure using a simulated microgravity rat model, with a focus on the pivotal role of acetazolamide (AZE) in regulating oculocranial biomechanical equilibrium. Experimental results demonstrated that short-term microgravity exposure elevated both IOP and intracranial pressure (ICP), while AZE intervention effectively mitigated these alterations, confirming its core mechanism of reducing aqueous humor production via carbonic anhydrase inhibition[46]. AZE likely alleviates microgravity-induced ocular stress through dual effects: IOP reduction and suppression of choroidal thickening. Mechanistically, AZE directly attenuates IOP elevation by inhibiting carbonic anhydrase in the ciliary epithelium, thereby reducing aqueous humor secretion[47]. Concurrently, its systemic action may indirectly modulate choroidal hemodynamics through ICP regulation. Previous studies indicate that AZE decreases cerebrospinal fluid (CSF) production rates[48], potentially alleviating optic nerve sheath pressure gradients and improving choroidal venous outflow resistance. These findings suggest that microgravity-associated fundus and choroidal congestion arises not only from intrinsic vascular dysregulation but also from impaired intracranial venous drainage [49]. Notably, this contrasts with the fluctuating IOP patterns observed during prolonged spaceflight: although short-term exposure induces marked IOP elevation, compensatory reductions in aqueous humor production during extended missions may restore IOP to baseline levels. Early AZE administration could proactively stabilize aqueous humor dynamics, preventing mechanical destabilization of the optic nerve head (ONH) biomechanical homeostasis, thereby offering a novel preventive strategy against spaceflight-induced visual impairment.

Prior research highlighting AZE’s inhibitory effects on choroidal thickness further reveals therapeutic potential beyond conventional IOP-lowering mechanisms. The choroidal vascular system, lacking robust autoregulatory capacity, undergoes volume expansion primarily governed by translamina cribrosa pressure gradients and hydrostatic forces[50]. Under microgravity-induced cephalad fluid shifts, elevated ICP may transmit retrograde via the optic nerve sheath, increasing choroidal venous pressure and vascular distension. Crucially, while retinal thickness remained stable during short-term exposure, rapid choroidal thickening in astronauts implicates hemodynamic perturbations from cephalad fluid redistribution as a key early pathogenic driver[51] [52]. Our data propose that AZE disrupts this vicious cycle through dual modulation of ICP and IOP: the AZE group exhibited significantly reduced translaminar cribriform pressure difference (TLCPD) compared to controls, with progressive attenuation of intraocular vascular mechanical strain over microgravity exposure. This aligns with clinical observations in ground-based microgravity analogs (e.g., head-down tilt bed rest), where AZE similarly demonstrates efficacy in stabilizing oculocranial pressure gradients [53] [54]

Minocycline, a neuroprotective tetracycline derivative, inhibits microglial and astrocytic activation [55]. Its efficacy has been extensively studied in vitro and across animal models of neurodegenerative diseases, including stroke, Alzheimer’s disease, Parkinson’s disease, multiple sclerosis, spinal cord injury, and traumatic brain injury. In retinal degenerative pathologies—such as branch retinal vein occlusion, ischemia-reperfusion injury, and light-induced retinal degeneration—minocycline exhibits anti-inflammatory and neuroprotective properties [56–58]. Microglia, resident immune sentinels in the central nervous system and retina, remain quiescent in healthy adults, localized perivascularly and within retinal layers[59]. Pathological stimuli (e.g., injury, pressure, or infection) trigger their proliferation, migration, and activation, often leading to redistribution within the optic nerve head [60, 61] Activated microglia release proinflammatory cytokines, reactive oxygen species, neurotoxic matrix metalloproteinases, and neurotrophic factors, while exhibiting phagocytic activity[62, 63]. Prior studies demonstrate that minocycline treatment suppresses excessive neuroinflammation by attenuating microglial and astrocytic activation, thereby preserving retinal integrity.

In chronic retinal degeneration models, minocycline effectively inhibits microglial activation and preserves visual function. To investigate whether retinal microglial inactivation protects against optic nerve degeneration during early glaucoma progression, we examined systemic, long-term minocycline administration in DBA/2J mice—a secondary glaucoma model replicating key human pathological features, including IOP elevation from anterior chamber drainage obstruction, IOP-dependent RGC axonal loss, gliosis, and microglial proliferation near RGCs [31, 64]. Our findings revealed that reduced microglial activation reversed the characteristic loss of RGC axonal transport and integrity typically associated with elevated IOP.In the present study, minocycline (MINO) significantly alleviated simulated microgravity-induced neurodegeneration by selectively suppressing aberrant microglial and astrocytic activation. Following short-term (2–4 weeks) simulated microgravity exposure, MINO intervention reduced expression of the microglial activation marker Iba-1 and reversed their morphology from pro-inflammatory “amoeboid” phenotypes to quiescent ramified states [64]. This effect correlated with MINO’s inhibition of the P2X7 receptor/NLRP3 inflammasome axis. Recent studies indicate that MINO suppresses inflammation by blocking NF-κB activation, attenuating LPS-induced microglial activation in vitro, and mitigating retinal degeneration in brain injury models[65, 66], including disease-associated microglia (DAM) [67]. Notably, studies have reported paradoxical exacerbation of visual loss and retinal degeneration under certain conditions. For instance, CX3CR1 signaling—mediated by neuron-derived CX3CL1—delivers “do not activate” signals to microglia [68, 69], while SOCS3 negatively regulates the JAK2/3-STAT3 pathway downstream of cytokine receptor signaling [70]. Future studies will explore whether minocycline modulates checkpoint proteins in cytokine receptor-activated pathways within microglia, potentially mitigating retinal toxicity.Our results align with prior findings in simulated microgravity hippocampal models, where MINO reversed neural stem cell (NSC) loss[71], underscoring its broad neuroprotective potential across diverse neurodegenerative contexts. Furthermore, by inhibiting hyperactivation of microglia and astrocytes, MINO markedly attenuated retinal ganglion cell (RGC) apoptosis and preserved visual function. Consistent with observations in chronic retinal degeneration models [56–58], these findings reinforce MINO’s therapeutic promise for retinal degenerative disorders In this study, we investigated two pharmacological interventions AZE and mino in a rat model of simulated microgravity. The treatments demonstrated varying degrees of visual protection. AZE, which modulates intraocular biomechanical pressure gradients, showed limited protective effects against short-term microgravity exposure. In contrast, minocycline, a neuroimmunomodulatory agent, significantly reduced retinal ganglion cell (RGC) apoptosis and preserved visual function, consistent with our previously proposed pressure-neuroimmunity theory.However, this study has several limitations. First, the simulated microgravity rat model requires refinement—for instance, adopting a centrifugation-based rotating platform could minimize tail-related mechanical stress and facilitate better animal monitoring. Second, both the microgravity exposure and drug intervention durations were relatively short. We hypothesize that under prolonged microgravity conditions, restoring pressure gradient homeostasis may become a critical strategy for visual protection. Additionally, the sample size was limited.Future work will focus on developing improved microgravity simulation methods, conducting comparative evaluations of different modeling approaches, and performing large-scale animal experiments to identify optimal protective interventions against simulated microgravity-induced visual impairment.

## Foundation

2021 National Natural Science Foundation of China project Personalized 3D Printed Physiological Myopia Correction Optical Lenses (52073181). No competing interests

## Conflicts of Interest

The authors declare no conflicts of interest.

## Notes

### Competing Interest Statement

The authors have declared no competing interest.

